# A screen for targets of the Drosophila pseudokinase Tribbles identifies Neuralized and Mindbomb, ubiquitin ligases that mediate Notch signaling

**DOI:** 10.1101/406249

**Authors:** Anna Shipman, Christopher Nauman, Britney Haymans, Rachel Silverstein, Leonard L. Dobens

## Abstract

Drosophila Tribbles (Trbl) is the founding member of a family of pseudokinases with conserved roles in antagonizing cell division, tissue growth and cell differentiation. In humans, three Tribbles isoforms serve as adaptor proteins, binding targets such as Cdc25 phosphatase, Akt kinase or the transcription factor C/EBP to block their activity or direct their proteosomal degradation. Mutations in Tribbles family members are associated with susceptibility to diabetes and cancer, notably Notch-induced tumor growth. Trbl misexpression in the fly wing disk leads to a block in mitosis associated with decreased levels of String/Cdc25 and increased levels of Cyclin B leading to reduced overall wing size and reduced trichome density. We show these Trbl growth-restricting phenotypes can be suppressed by manipulating levels of known Trbl targets, and use this sensitized wing system to screen a collection of growth regulating open reading frames (ORFs) to search for enhancers and suppressors affecting cell and tissue size. By precisely measuring morphometric changes in wing phenotypes using a computer-based tool, we detected synthetic interactions with several E3 ubiquitin ligases, and focused our analysis on the Notch pathway components Neuralized (Neur) and Mindbomb1 (Mib1). In the wing, notum and egg chamber epithelia, Trbl misexpression suppressed Neur and Mib1 activities and stabilized the accumulation of both proteins. To understand these interactions, we used yeast two-hybrid assays to show Trbl physically bound to both Neur and Mib1. Our data are consistent with published reports that mammalian Tribbles3 modulates Notch responses by binding and stabilizing Mindbomb and indicate that a wing misexpression approach is useful to identify novel components in a conserved Tribbles signaling pathway.

**AUTHOR SUMMARY:** Tribbles pseudokinases are adaptor molecules, binding diverse targets regulating cell differentiation, growth and proliferation and directing their proteasomal degradation. To search for novel targets of Drosophila Tribbles, we adopted a wing co-misexpression scheme and measured changes in cell/tissue size to identify enhancers and suppressors of the Tribbles phenotype. We show the Notch pathway components Neuralized and Mindbomb1 E3 ligases act as Tribbles suppressors and demonstrate that Tribbles modulates their levels and activites. Recent demonstration that mammalian Tribbles 3 binds the E3 ligase Mindbomb to promote ligand-mediated Notch activation implies a conserved role for Tribbles family members in Notch activation.

## INTRODUCTION

Several unrelated screens for Drosophila genes required for cell division and cell migration conducted in the year 2000 converged on the gene Tribbles (Trbl), an adaptor protein that facilitates the proteosomal turnover of key regulators of cell differentiation (Grosshans and Wieschaus, 2000; Rorth et al., 2000; Mata et al., 2000; Seher and Leptin, 2000). Subsequent unbiased fly genetic screens using both misexpression and loss-of-function approaches have uncovered diverse roles for Trbl in (1) neural differentiation during bristle patterning (Abdelilah-Seyfried et al., 2000), (2) stem cell proliferation in the germline (Schulz et al., 2004), (3) JAK/STAT signaling to modulate eye growth (Mukherjee et al., 2006), and (4) memory formation in the adult brain (LaFerriere et al., 2008).

Trbl belongs to a family of proteins that share a central DLK catalytic loop motif characteristic of canonical kinases, however the absence of both ATP and Mg2+ binding motifs typical of bona fide kinases has placed these genes in the category of pseudokinases (Wilkin et al., 1996; Wilkin et al., 1997; Mayumi-Matsuda et al., 1999). At the C-terminus of Trib family members, conserved COP1 and MEK1 binding sites interact with targets including MAPK kinases, transcription factors, membrane receptors and E3 ligases to alternatively degrade, stabilize or block target activation, depending on the tissue examined (reviewed in Sakai et al., 2016; Ord, 2017; Eyers et al., 2017). By linking diverse targets to the proteosomal machinery, Tribbles family members act as tumor suppressors or oncogenes depending on cellular context, and variants have been connected to diseases such as diabetes (Nakamura, 2015; Salazar et al., 2015; Stein et al., 2015).

The list of known Drosophila Trbl targets is short but highly conserved with mammalian targets, including the Cdc25 phosphatases String and Twine (Mata et al., 2000; Farrell and O’Farrell, 2013; Liang et al., 2016b), the C/EBP homolog Slbo (Rorth et al., 2000; Wouters et al., 2007; Dedhia et al., 2010; Keeshan et al., 2010; Masoner et al., 2013), and Akt kinase (Du et al., 2003; Das et al., 2014; Salazar et al., 2015; Fischer et al., 2017). To identify novel components acting in the Trbl signaling pathway, we conducted a wing misexpression screen of 663 transgenic fly lines, each harboring an open reading frame (ORF) encoding a growth regulatory gene, in a Trbl-sensitized background (Schertel et al., 2013). We used the ImageJ-based tool Fijiwings (Dobens and Dobens, 2013) to evaluate wing phenotypes and from this approach identified both enhancers and suppressors of Trbl misexpression phenotypes. Among Trbl interactors, we identified the E3 ubiquitin ligases Neuralized (Neur) and Mindbomb (Mib1), and several tests of their interactions with Trbl indicate a conserved role in modulating Notch responses.

## RESULTS

### The Trbl wing misexpression phenotype shows interactions with known targets

The Drosophila wing disk primordium is an epithelial sheet that proliferates, grows in size and is patterned along the anterior-posterior and proximal distal axes during larval development (Fig. 1A). The wing blade evaginates at late metamorphosis (Fig. 1B) and the dorsal and ventral layers of the wing blade adhere to secrete a cuticle that hardens into a mature wing marked by a characteristic vein pattern (Fig. 1C). The wing is covered by non-neuronal trichome structures, which arise from actin-rich extensions produced distally from each cell in the wing epithelial bilayer.

**Fig. 1.**
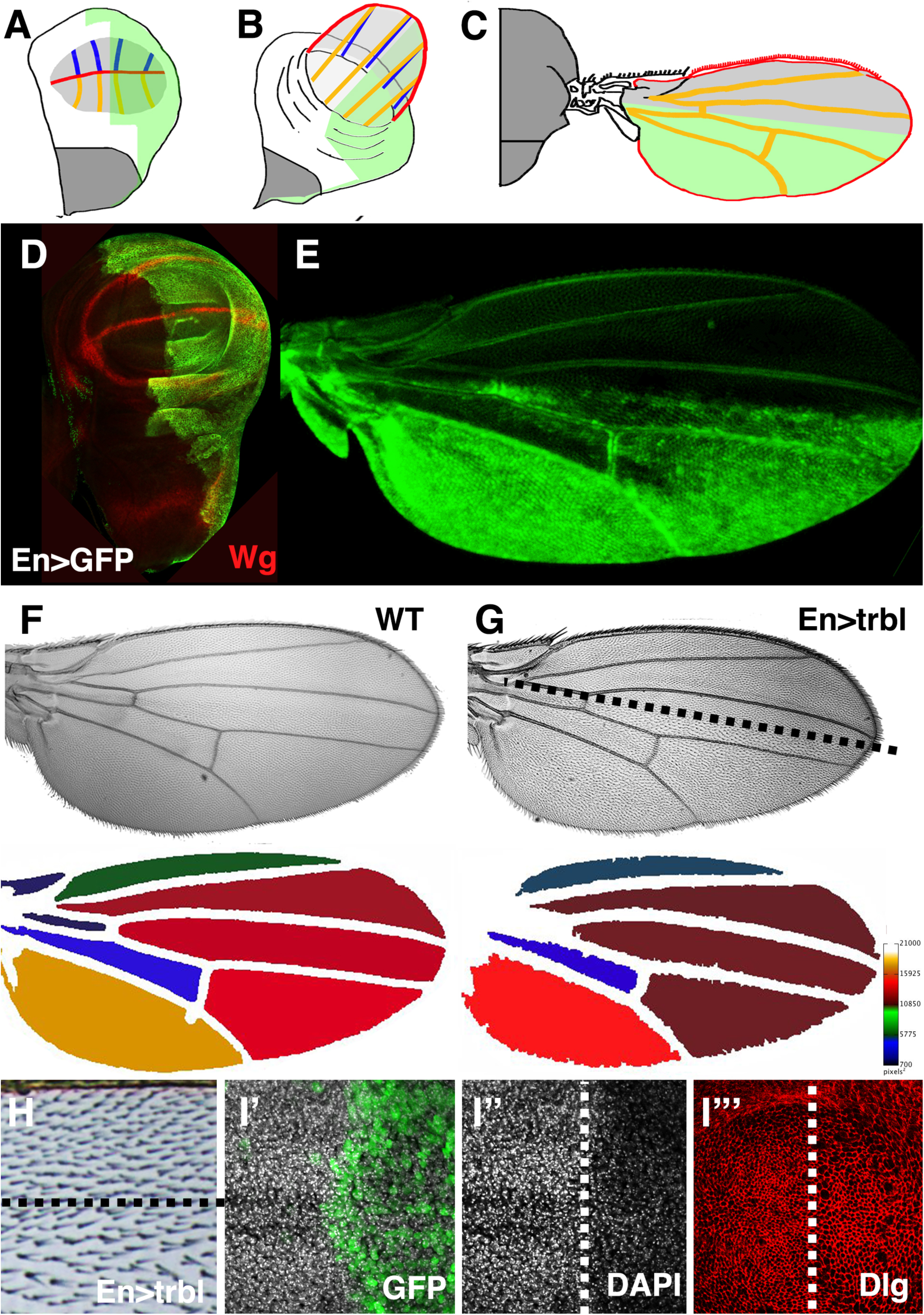
Wing cell size and tissue size are sensitive to Trbl misexpression. (A-C) Representation of EngrailedGAL4 (EnGAL4) expression (green) in the posterior compartment of the everting wing disk where the wing blade primordia are in light gray, the notum in dark gray, the forming wing margin in red, the dorsal wing veins in orange and the ventral wing veins in blue. (D, E) EnGAL4, UAS-GFP expression (green) in the posterior compartment of the wing disk (green, A) where expression of Wingless (Wg, red) marks the wing margin and hinge domain. In the mature wing prior to cuticular hardening, GFP expression can be detected in the posterior compartment (E). (F) Wild type wing (WT, top) with heat map (below) showing relative size of intervein regions based on kilopixels of each region. (G,H) EnGAL4 driving misexpression of UAS-Trbl in mature wing resulted in a reduction in the density of trichomes in posterior compartment compared to the anterior compartment where UAS-trbl is not expressed. The compartment boundary is represented by dotted line. Heat map (below, G) shows reduction in overall wing size compared to WT wing. (I-I’”) Wing disk misexpressing UAS-Trbl and UAS-GFP (green) in posterior wing compartment (right side where boundary is represented by dotted line) revealed reduced cell number reflected as reduced DAPI nuclear staining (I”) and increased cell size whose borders are outlined by Discs large (Dlg, red) accumulation, I”’.

Engrailed-GAL4 (En-GAL4) driving UAS-GFP expression marks the posterior compartment of the wing disk primordium and the corresponding posterior region of the mature wing (Fig. 1D,E; Yoffe et al., 1995). Misexpression of Trbl in this domain has two effects on wing morphology (noted in Dobens and Dobens, 2013; Mata et al., 2000): first, the overall size of the wing is reduced compared wild type (WT; Fig. 1F,G); and second, trichomes are less dense than in the anterior half (Fig.1H). An increase in wing cell size was seen with misexpression of one copy of Trbl (Fig. 1I) and consistent with a block in cell division at G2, we observed increased levels of Cyclin B (CycB) in the posterior of these wing disks, detected both directly by antisera (Fig. 2G) and indirectly by a slight stabilization of a ubiquitously expressed RFP-CycB transgene (Supplemental Fig. 1; Zielke et al., 2014). The wing is sensitive to further increased levels of Trbl, so that EnGAL4 driving two copies of Trbl is adult lethal (Fig. 2A,D).

**Fig. 2.**
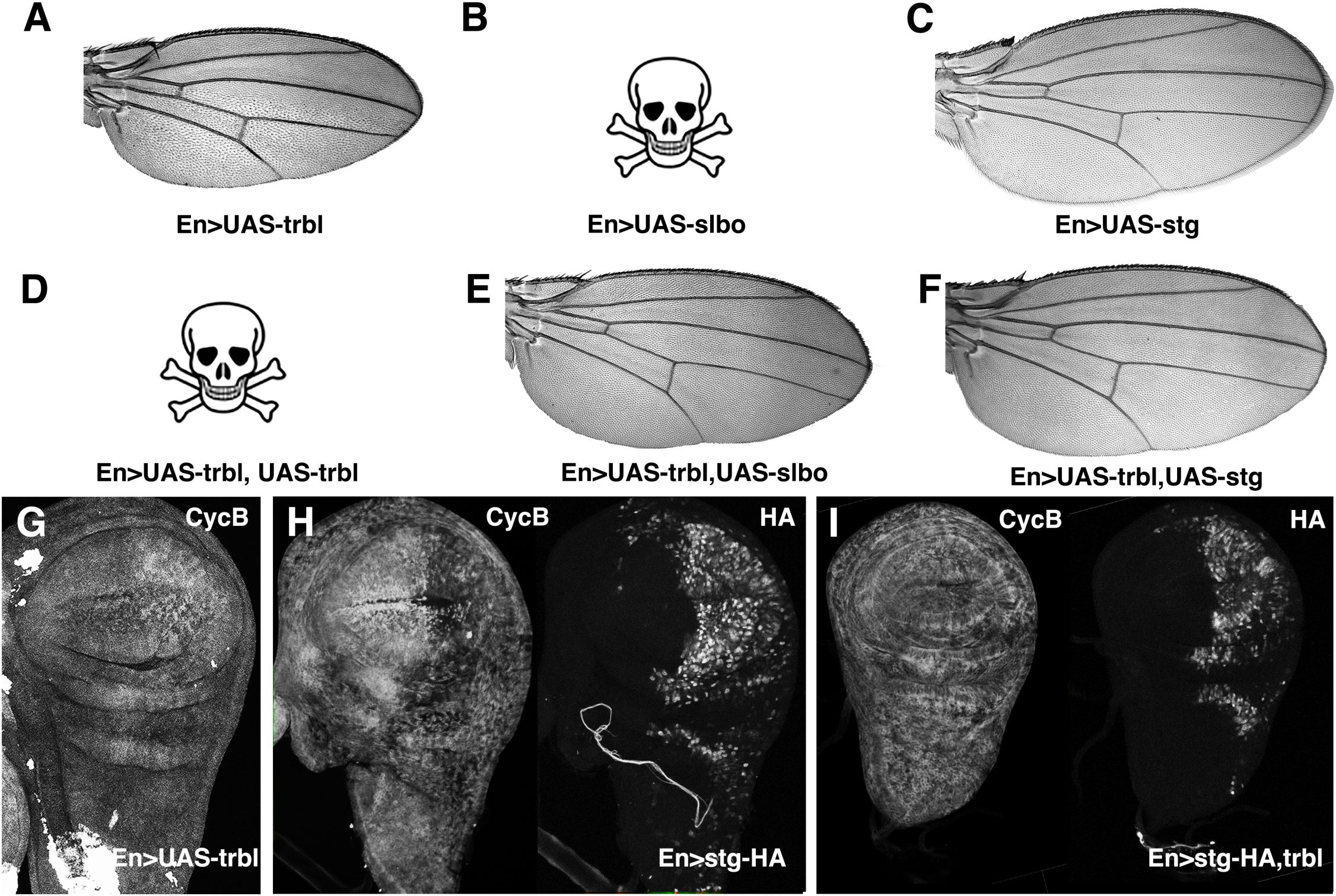
The Trbl targets Stg and Slbo suppress Trbl wing phenotypes. A. Wing resulting from EnGAL4 driving misexpression of UAS-trbl. B. EnGAL4 driving misexpression of UAS-slbo was lethal. C. Wing resulting from EnGAL4 driving misexpression of UAS-string. D. EnGAL4 driving misexpression of two copies of UAS-trbl was lethal. E. A normally patterned wing resulting from EnGAL4 driving misexpression of UAS-slbo and UAS-trbl in rare escapers. F. Wing resulting from EnGAL4 driving misexpression of UAS-string and UAS-trbl has reduced cell density compared to UAS-Trbl alone (A). G. Cyclin B (CycB) accumulation was increased when UAS-trbl is misexpressed in the posterior compartment by EnGAL4. H. Cyclin B accumulation (left panel) was decreased when UAS-HA-string is misexpressed at high levels (right panel) in posterior compartment by EnGAL4. I. Cyclin B accumulation (left panel) was comparable to WT levels when UAS-HA-string is co-misexpressed with UAS-trbl (right panel) in posterior compartment by EnGAL4. HA-Stg levels are reduced as well (Cf. H.I, right panels for each).

To test if these wing phenotypes were sensitive to interactions with known Trbl targets, we first examined the C/EBP encoded by the *slow border cells* (*slbo*) gene (Montell et al., 1992; Rorth et al., 2000). EnGAL4 misexpressing UAS-slbo was lethal, likely due to its potent transcription factor activity (Fig. 2B). Trbl co-expression with Slbo weakly rescued this lethality, resulting in rare escapers with well-patterned wings (Fig. 2E), consistent with the ability of Trbl to bind and degrade Slbo protein when co-expressed (Rorth et al., 2000). Next we examined the interaction between Trbl and its target String (Edgar and O’Farrell, 1990; Grosshans and Wieschaus, 2000; Mata et al., 2000). EnGAL4 misexpression of an HA-tagged version of String in the wing disk led to a clear decrease in CycB protein levels (Fig. 2H) and destabilization of a CycB-RFP fusion (Supplemental Fig. 1B) in the posterior region, an effect opposite to that of Trbl on CycB levels (cf. Fig. 2 G,H) and consistent with the ability of increased String to promote cell division in this region. When HA-tagged String was co-expressed with Trbl in the wing disk we observed: (1) suppression of the Trbl wing phenotype, so that mature wing size was normal and the density of trichomes between anterior and posterior compartments was similar (Fig. 2C,F); (2) levels of CycB in the posterior wing disk tissue that were comparable to levels in the anterior (Fig. 2I), and (3) reduced levels of HA-String compared to misexpression of HA-Stg alone (cf. 2H,I). These observations are consistent with the demonstrated ability of Trbl to direct Stg turnover to block cell division (Grosshans and Wieschaus, 2000; Mata et al., 2000). Based on these Trbl-String and Trbl-Slbo interactions we conclude that the wing is an effective place to search for Trbl target genes, and that the HA-tag can reveal the effects of Trbl on candidate target stability.

### Screen for enhancers and suppressors of Trbl cell and tissue size phenotypes

Based on Trbl interactions with its known targets, we conducted a large-scale screen for novel Trbl interactors using a set of strains bearing UAS-regulated open reading frames (ORF) focusing on components from the insulin, wingless/Wnt, Notch and other signaling pathways known to regulate cell division and growth (Schertel et al., 2013). We crossed a strain bearing both the EnGAL4 driver and UAS-trbl to this UAS-ORF collection (Fig. 3A) and prepared wings from F1 progeny co-misexpressing both transgenes.

**Fig. 3.**
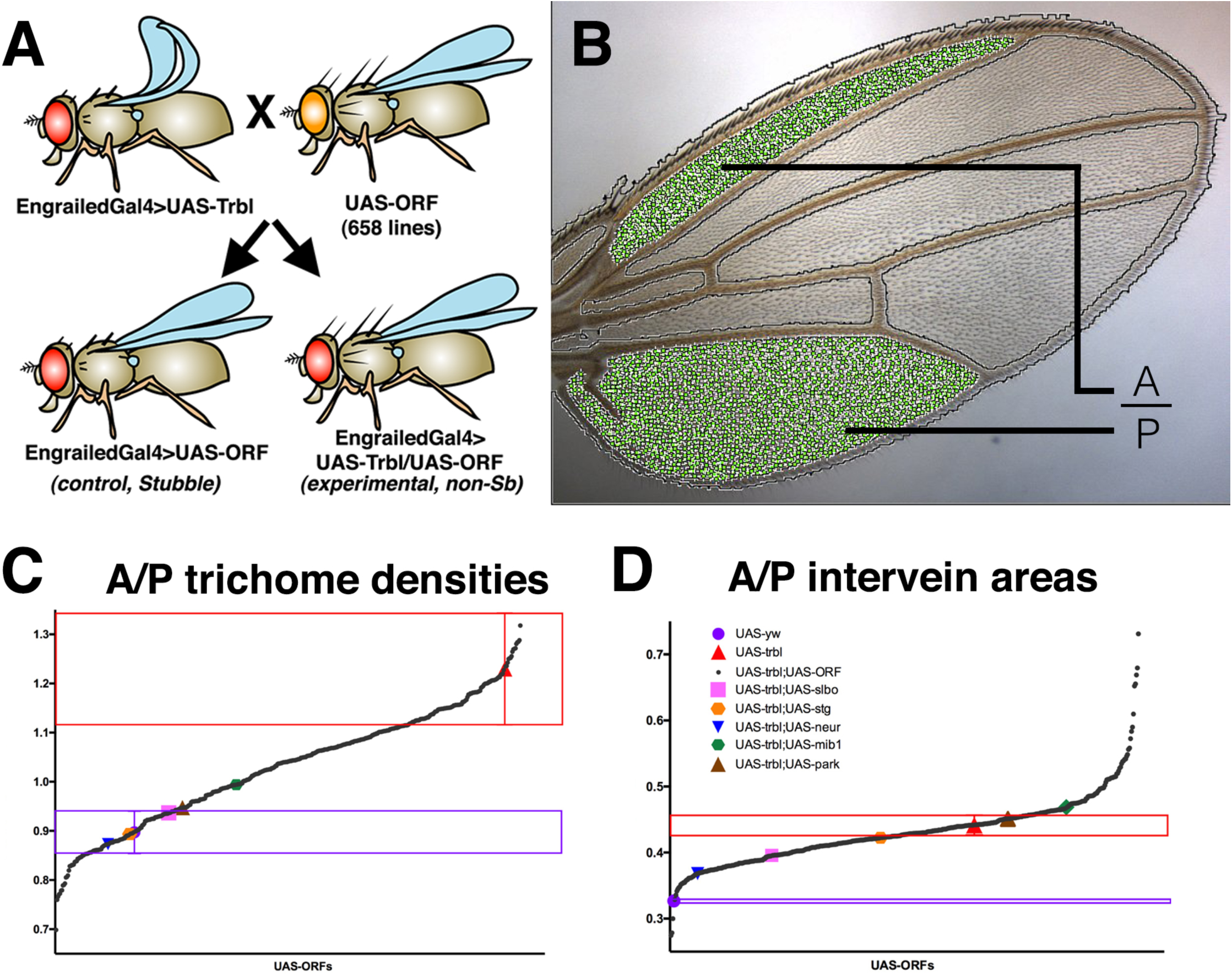
A genetic screen for Trbl modifiers of wing cell and tissue size. A. Genetic scheme to detect Trbl misexpression interactors. Females bearing the EnGAL4 driver and UAS-trbl transgene were crossed to males from 663 strains bearing different UAS-ORFs and F1 progeny were identified based on cuticle markers. B. Wings from flies misexpressing UAS-trbl and a UAS-ORF were analyzed using Fijiwings (Dobens and Dobens, 2013) for size and trichome density specifically in the anterior-most and posterior-most intervein regions and the ratio of these values calculated for (C) and (D). C. The average anterior/posterior trichome density ratio of all UAS-ORFs co-misexpressed with Trbl by the EnGAL4 driver, in order from lowest to highest ratio. Highlighted at the two extremes are the UAS-yw (WT) trichome density ratio value in purple and the Trbl trichome density ratio value in red. The red box centered the Trbl red dot is designed to define a range of cell size ratios deemed not significantly different from the Trbl phenotype. ORFs that fall within the purple box represent the range of measured ratios within one standard deviation of WT and include the strong suppressors Stg and Slbo, highlighted in orange and pink, respectively. Weaker Trbl phenotype suppressors fall in between the WT and Trbl ranges. D. The average anterior/posterior area ratio of all UAS-ORFs co-misexpressed with Trbl by the EnGAL4 driver, in order from lowest ratio to highest ratio. The red box and purple line represent one standard deviation from the average area ratio of UAS-trbl and UAS-yw respectively. Highlighted at the two extremes are the UAS-yw (WT) intervein area ratio value in purple and the Trbl ratio value in red. The purple box is centered on the range within one standard deviation of a UAS-yw WT ratio while the red box represents the range not significantly different from the area ratio of a UAS-trbl wing. Co-expressed ORFs within the red box are deemed without effect on the Trbl misexpression area phenotype, strong suppressors would be within the purple WT box, and weak suppressors fall between these ranges.

In order to quantify statistically significant changes to tissue size and cell number caused by ORF/Trbl co-misexpression, we used Fijiwings (Dobens and Dobens, 2013), a set of macros applied to the FIJI build of ImageJ allowing semi-automated wing measurements of (1) trichome density, which as noted above correlates with cell number, and (2) intervein region size, a proxy for tissue size (Fig. 3B). Because EnGAL4 is expressed in the posterior compartment while the anterior compartment is unaffected, we calculated the ratio of trichome density of the first anterior and third posterior intervein regions and separately calculated the ratio of areas between of the first anterior and third posterior intervein regions (Fig. 3B). In this way, we corrected for the influence of nutrition, crowding and environment, which affects the measurements of cell and tissue size in both domains similarly (Table 1).

From the screen we obtained three categories of interactions (Tables 2 and 3). First, ORFs whose lethality was suppressed by Trbl (an example is Slbo, detailed above). This class included the C2H2 zinc finger transcription factor Castor (Cas; Mellerick et al., 1992). Second, we identified ORFs with subtle defects in cell and tissue size that were suppressed by Trbl, a class of interactors reminiscent of wing interactions phenotypes seen with Stg (Fig. 2C,F). This class included (1) the BTB zinc finger transcription factor Lola-like (Lolal; Giniger et al., 1994; Quijano et al., 2016) (2) the CCR4-NOT transcription complex subunit Not3 (Temme et al., 2004), and (3) the E3 ubiquitin ligase Parkin (Park; Yang et al., 2003). And finally, we identified ORFs whose misexpression resulting in strong wing patterning phenotypes that were effectively suppressed by Trbl; this class included (1) the WD repeat protein Will die slowly (Wds; Hollmann et al., 2002), (2) the Wingless homolog Wnt oncogene analog 5 (Wnt5 a.k.a Wnt3; Russell et al., 1992) and (3) the E3 ubiquitin ligase Neuralized (Neur, detailed below; Boulianne et al., 1991).

In Fig. 3C, average trichome density ratio values derived from measurements of co-misexpression of Trbl and each one of the UAS-ORF genes tested in the screen are ordered from lowest to highest, and in Fig. 3D wing area measurements ratios are summarized ordered from lowest to highest. For both trichome density and wing area, strong suppressors of the Trbl phenotype fall close to WT wing measurements while weak suppressors fall between the Trbl-WT extremes. Notably, a few potential enhancers of the Trbl area phenotype were detected, exhibiting even higher area ratios when co-misexpressed.

When we simultaneously compared the measures of tissue size and cell size using a scatter plot (Supplemental Fig. 2), the relatively strong increase in cell size paired with a decrease in tissue size characteristic of Trbl misexpression placed its combined phenotypes at the far upper right extreme of measurements (shown as a circled red triangle), while WT wings occupied the bottom left extreme of the plot (circled purple circle, left). Co-misexpression of most ORFs suppressed both Trbl phenotypes, resulting in a continuous “line” of suppression from right to left. Notably the Trbl suppressors Stg and Slbo, highlighted in orange and pink respectively, were located near the WT point, consistent with their strong suppression of both aspects of the Trbl phenotype. Notably, misexpression of some ORFs led to larger tissue size and in some cases much smaller tissue size, placing these interactions above or below the continuum and pointing to genes with differential influences on Trbl regulation of cell size vs. cell number.

In summary, Fijiwings analysis allowed the detection of subtle differences in cell and tissue size to identify weak suppressors and enhancers that potentially interact with Trbl both directly and indirectly, in pathways upstream, downstream or parallel to Trbl.

### Trbl modulates the strength of Notch signaling by stabilizing Neuralized and Mindbomb1 and suppressing their activity

Among the Trbl genetic suppressors in our UAS-ORF screen were several E3 ubiquitin ligases, including Wds, Neur, and Parkin (the latter two highlighted in blue and brown, respectively, in Fig. 3B,C and Supplemental Fig. 2). Consistent with this class of interactors, human Trib1 has been demonstrated to directly bind with high affinity the E3 ligase COP1 (Uljon et al., 2016). Recent genome-wide screens have identified a Neuralized binding site in Trbl (data not shown and Dinkel et al., 2016), so we focused on this E3 ligase and tested its interactions with Trbl in more detail.

Trbl misexpression strongly suppressed Neur misexpression wing notch phenotypes (cf. Fig 4A,B) and conversely Neur suppressed strongly the Trbl large cell phenotype (Fig. 4I). WT Trbl effectively suppressed the split wing phenotype associated with misexpression of a Neur transgene deleted for the RING domain, so Trbl-Neur suppression was not dependent on this conserved motif (Fig. 4C,D).

**Fig. 4.**
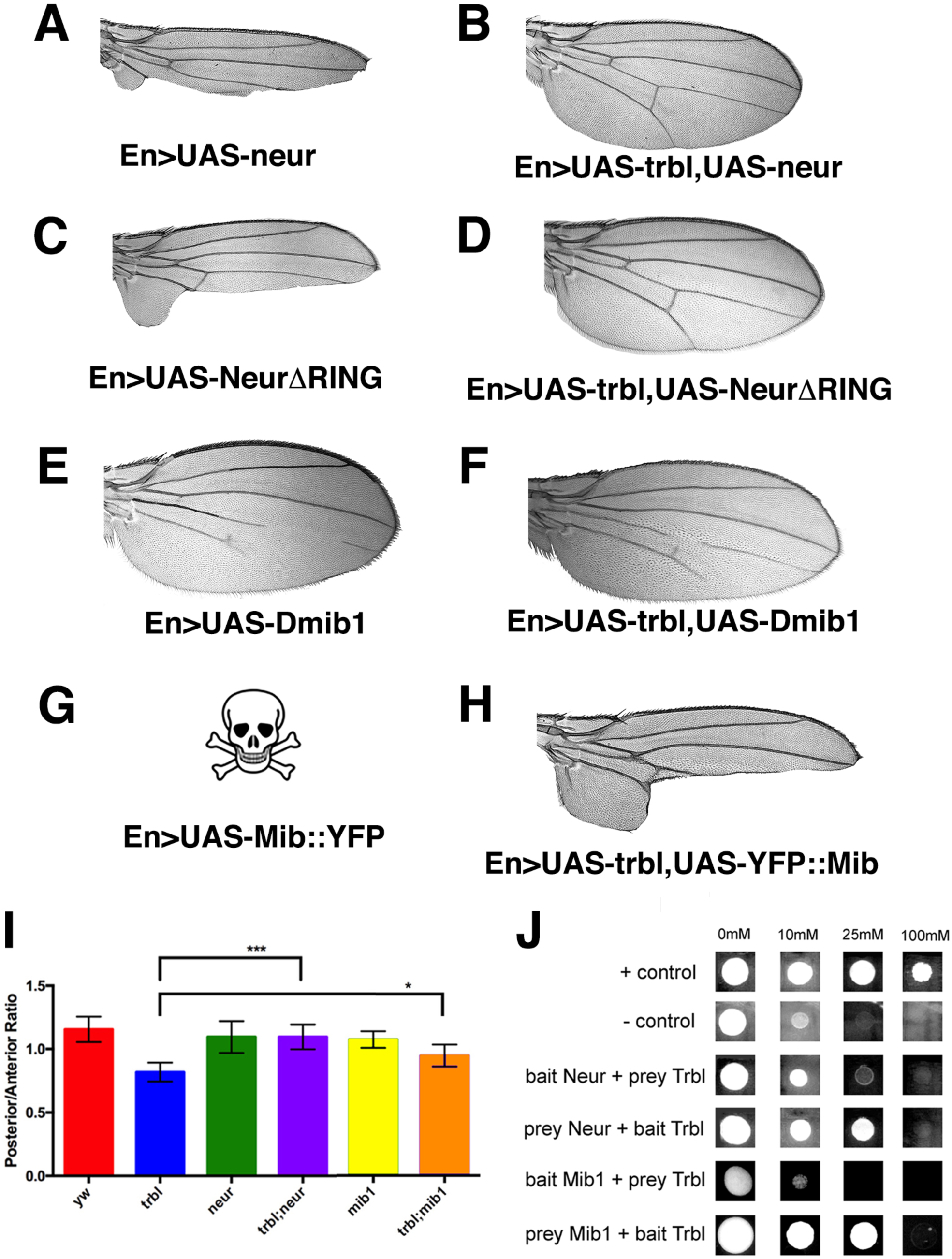
Trbl suppresses Neuralized and Mindbomb to promote Notch signaling. A. Wings from flies misexpressing UAS-neur under the influence of the EnGAL4 driver showed wing notches in the posterior compartment. B. Wings from flies misexpressing UAS-neur and UAS-trbl under the influence of the EnGAL4 driver suppressed the wing notch phenotypes seen in (A). C. Wings from flies misexpressing UAS-neurΔRING under the influence of the EnGAL4 driver showed split wing phenotype in the posterior compartment. D. Wings from flies misexpressing UAS-neurΔRING and UAS-trbl under the influence of the EnGAL4 driver suppressed the split wing phenotypes seen in (C). E. Wings from flies misexpressing EnGAL4; UAS-mib1 showed wing vein defects in posterior compartment. F. Wings from flies misexpressing UAS-mib1 and UAS-trbl under the influence of the EnGAL4 driver suppressed the wing vein phenotypes seen in (E). G. EnGAL4 misexpression of a YFP-tagged mib1 transgene (UAS-Mib::YFP) resulted in F1 lethality. H. Wings from flies misexpressing UAS-Mib::YFP and UAS-trbl under the influence of the EnGAL4 driver suppressed the lethality of UAS-Mib::YFP alone and lead to a split wing phenotypes distinct from the notch phenotype in (A). I. Trichome density ratios compared between UAS-yw, UAS-trbl, UAS-neur; UAS-trbl+UAS-neur, UAS-mib1 and UAS-trbl;UAS-mib1. Note that UAS-YFPMib1 misexpression in the wing alone was lethal. An unpaired T-test between Trbl and Trbl; Neur wings had a p-value of 0.0006, indicated on the graph by ***. An unpaired t test between Trbl and Trbl; Mib1 wings had a P-value of 0.0371, indicated by *. The error bars represented one standard deviation from the mean. N = 8 for UAS-yw, N = 4 for UAS-trbl, N = 4 for UAS-neur, N = 8 for UAS-trbl+UAS-neur, N = 3 for UAS-mib1, and N = 7 for UAS-trbl;UAS-mib1. J. Yeast transformed with Neur or Mib1 (as prey) and Trbl (as bait) is able to grow on plates containing up to 25mM 3-AT, indicating a weak interaction. Yeast transformed with Neur or Mib1 (as bait) and Trbl (as prey) showed a weaker interaction (up to 10mM 3-AT), no stronger than a negative control. The positive and negative controls are described in the Materials and Methods.

To understand the mechanism for this effect of Trbl on Neur misexpression phenotypes, we tested for physical interactions with Neur using the yeast two hybrid assay (Fields and Song, 1989). In silico evaluation of Trbl detects a Neur binding site (Dinkel et al., 2012; Fazekas et al., 2013) and consistent with this, we detected weak Trbl-Neur binding under selective conditions (up to 25mM 3AT) in the Neur prey/Trbl bait configuration (Fig. 4J). Because Trbl suppresses Neur wing margin phenotypes, we examined the effect of co-misexpressing Trbl and HA-Neur on Neur stability and the expression of the margin-specific Notch target gene Wingless (Wg). Misexpression of HA-Neur alone in the posterior wing blade under the influence of EnGAL4 is sufficient to reduce expression of Wg specifically in the posterior domain (Fig. 5A, left panel; Brook and Cohen, 1996), consistent with reduced tissue growth at the margin that results in the wing notch phenotype observed (Fig. 4A). EnGAL4 co-misexpression of Trbl with HA-Neuralized in the posterior wing suppressed the Neur wing notch phenotype (Fig.4B) and consistent with this, we observed increased Wg expression, associated with tissue growth and patterning, in the wing margin domain (arrow, Fig. 5B left panel).

**Fig. 5.**
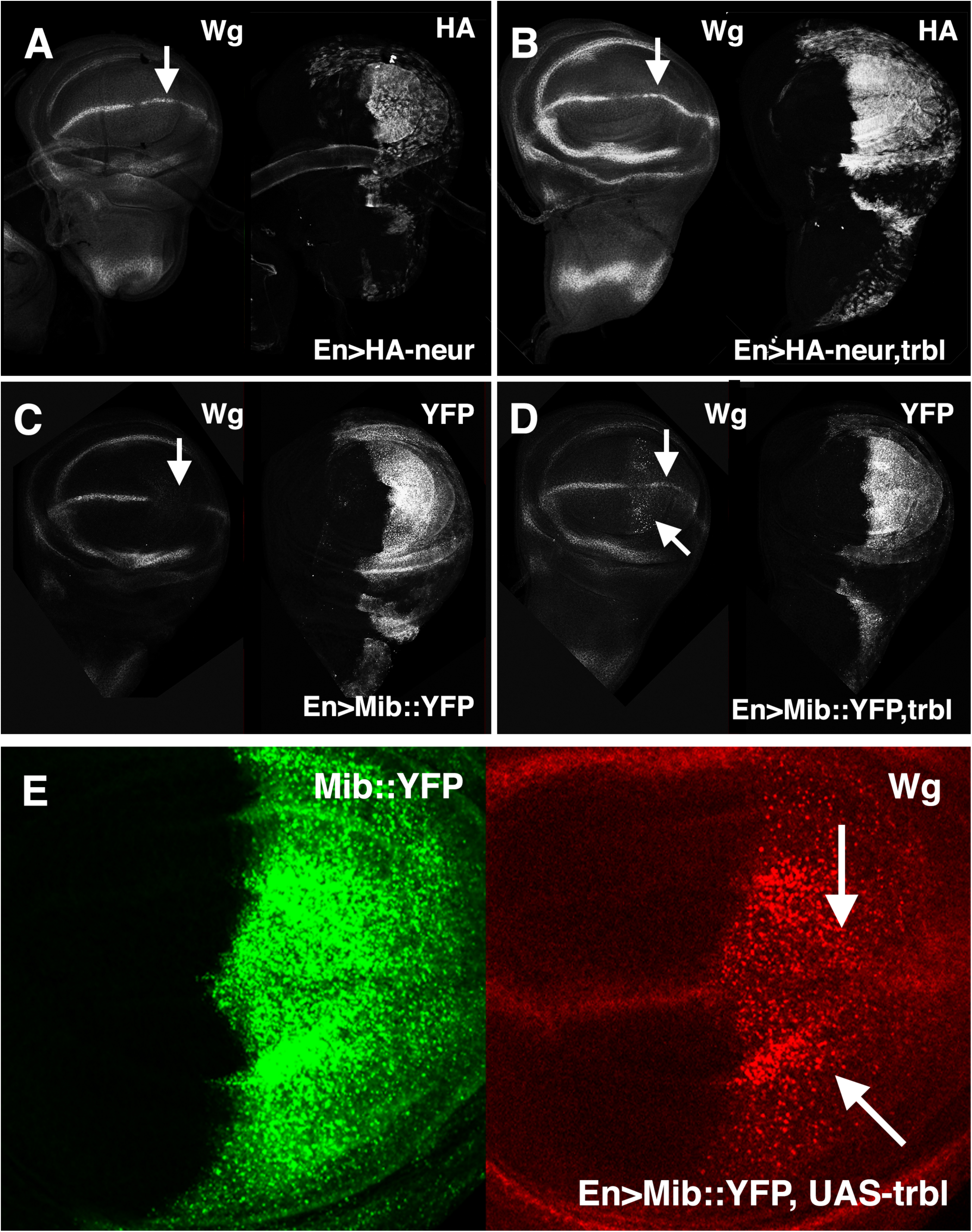
Trbl interacts with Neuralized and Mindbomb to pattern the wing margin. A. EnGAL4 misexpression of UAS-HA-neur as detected by HA antisera (right panel) reduced expression of Wingless (Wg, left panel, arrow) in the posterior compartment of the wing. B. EnGAL4 misexpression of UAS-HA-Neur and UAS-trbl restored strong expression of Wingless (Wg, left panel, arrow) to the posterior compartment of the wing and led to increased HA-neur levels. C. EnGAL4 misexpression of UAS-Mib1::YFP, a YFP-tagged version of Mindbomb (right panel) strongly reduced expression of Wingless (Wg, left panel, arrow) in the posterior compartment of the wing. D. EnGAL4 misexpression of UAS-Mib1::YFP and UAS-trbl restored strong expression of Wingless (Wg, left panel, arrow) to the posterior compartment of the wing and led to no strong effect on Mib1::YFP levels. E. Closeup of wing blade from EnGAL4; UAS-Mib1::YFP /UAS-trbl wing disk similar to (D) shows strong ectopic expression of Wg both at that posterior margin and in flanking cells (arrows) consistent with Trbl activation of Mib-mediated Notch signaling in this domain.

Because Trbl destabilized an HA-tagged version of Stg to block its activity in the wing (Fig. 2H), we examined the effect of co-misexpressing Trbl and HA-tagged Neur on HA-Neur levels. In contrast to its effect on HA-Stg, co-misexpression of Trbl and HA-Neur resulted in an increase in HA levels (Fig. 5B, right panel) suggesting that Trbl stabilizes Neur levels in the wing primordium.

To test Neur and Trbl interactions in tissues where they both are normally expressed, we used Neuralized-GAL4 (NeurGAL4), which is expressed in the scutellar sensory organ precursor cells (revealed by UAS-GFP, Fig. 6A and Supplemental Fig. 3B) that form the scutellar bristles on the mature notum (Fig. 6B-D). Stg misexpression by NeurGAL4 leads to loss of bristles, likely due to disruption of the ordered division of sensor organ precursor (SOP) cells that pattern these bristles (Fig. 6E) while co-expression of Trbl effectively suppresses the effect of Stg (Fig. 6F) indicating that Trbl can block its target Stg in this tissue in a manner similar to the wing. It has been shown that NeurGAL4 misexpression of Trbl leads to loss of bristles, a phenotype similar to the effect of NeurGAL4 driving RNAi knockdown of neuralized (Supplemental Fig. 2 and Le Borgne and Schweisguth, 2003; Rhyu et al., 1994; Weinmaster, 1998). As shown in Fig. 6G and H, bristle loss after NeurGAL4 misexpression of Neur is suppressed effectively by co-expression of Trbl (cf. Fig. 6G,H), suggesting that Trbl blocks Neur activity to coordinate the ordered cell division and cell fate decisions that pattern these bristles.

**Fig. 6.**
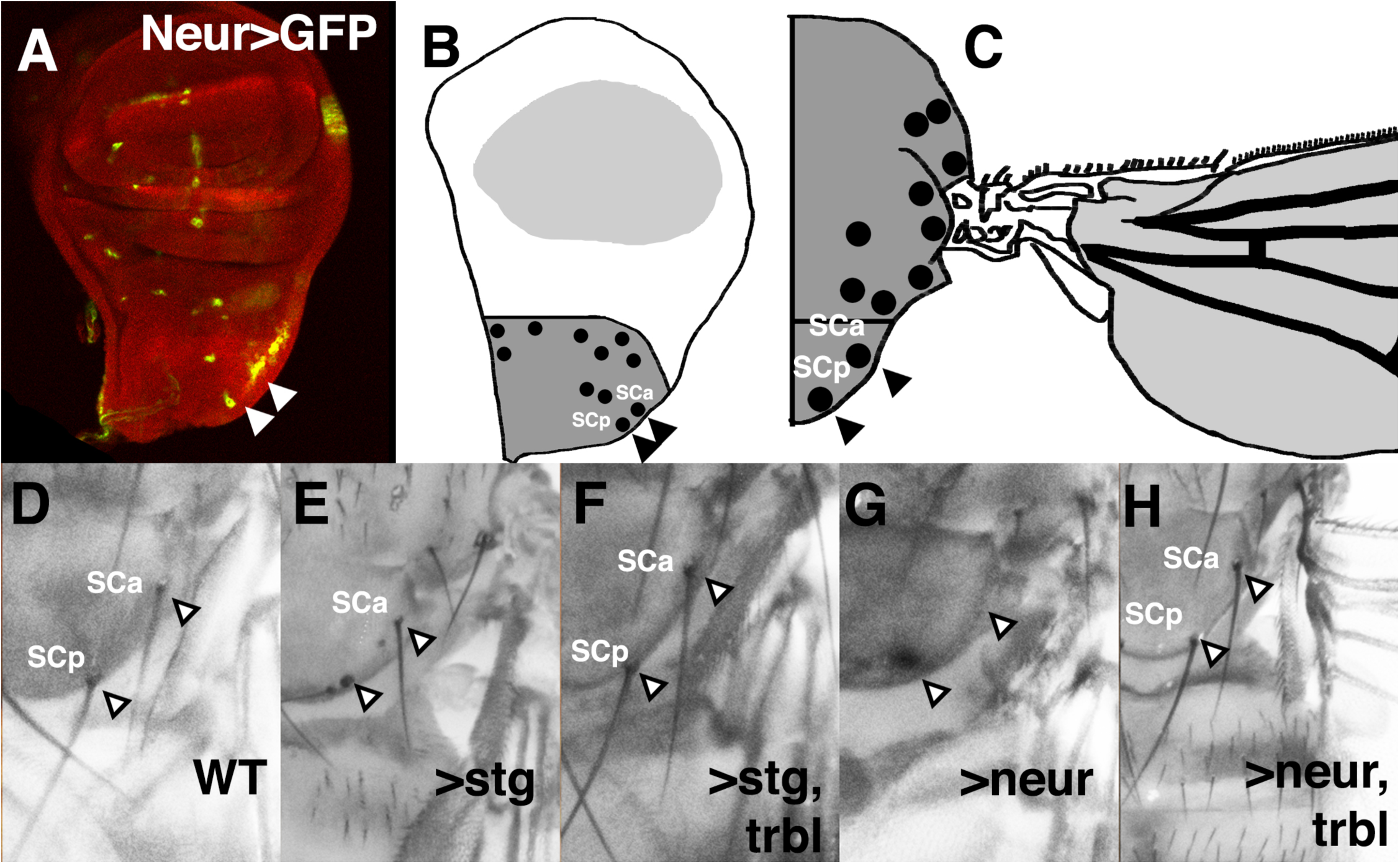
Trbl interacts with Neuralized during scutellar bristle patterning. A-C. Wing disk misexpressing NeurGAL4;UAS-GFP (green, A) marks the developing sensory organ precursor cells in the notum, represented in (B) as notal bristle SOPs (black dots) in notal primordium (dark gray), which differentiate as anterior scutellar bristle (SCa) and posterior scutellar bristle (SCp) primordia on corresponding mature notum (C). D. Arrowheads point to the position of anterior scutellar bristle (SCa) and posterior scutellar bristle (SCp) in wild type (WT) notum. E. Posterior scutellar bristles were missing in NeurGAL4 driving UAS-string. F. Posterior scutellar bristles were restored in NeurGAL4 driving UAS-string and UAS-trbl, indicating suppression of the Stg phenotype in (E). G. Anterior and posterior scutellar bristles were missing in NeurGAL4 driving UAS-neur. H. Bristles were restored in NeurGAL4 driving UAS-neur and UAS-trbl, indicating suppression of the Neur phenotype in (G).

Next, we examined the ovarian border cells, where Trbl and Neur are also both expressed (data not shown and Masoner et al., 2013). First, we showed that HA-Stg misexpressed by the Slow border cells-GAL4 (SlboGAL4) driver (Rorth et al., 1998) resulted in high levels of HA staining that were reduced to nearly undetectable levels when co-expressed with Trbl (cf. Fig. 7A,B). This indicates that as in the wing, Trbl effectively turns over Stg protein in the border cells. In contrast to its effects on Stg, SlboGAL4 co-misexpression of Trbl and HA-Neur effectively increased levels HA-Neur in the migrating border cells, compared to SlboGAL4 misexpression of HA-Neur (Fig. 7C,D). Similar stabilization of Neur with Trbl was observed with another border cell driver GAL4-Hsp70.PB (Supplemental Fig. 4). Together, these data show that Trbl can selectively destabilize targets like String while stabilizing other targets like Slbo and Neuralized with consequences for cell differentiation.

**Fig. 7.**
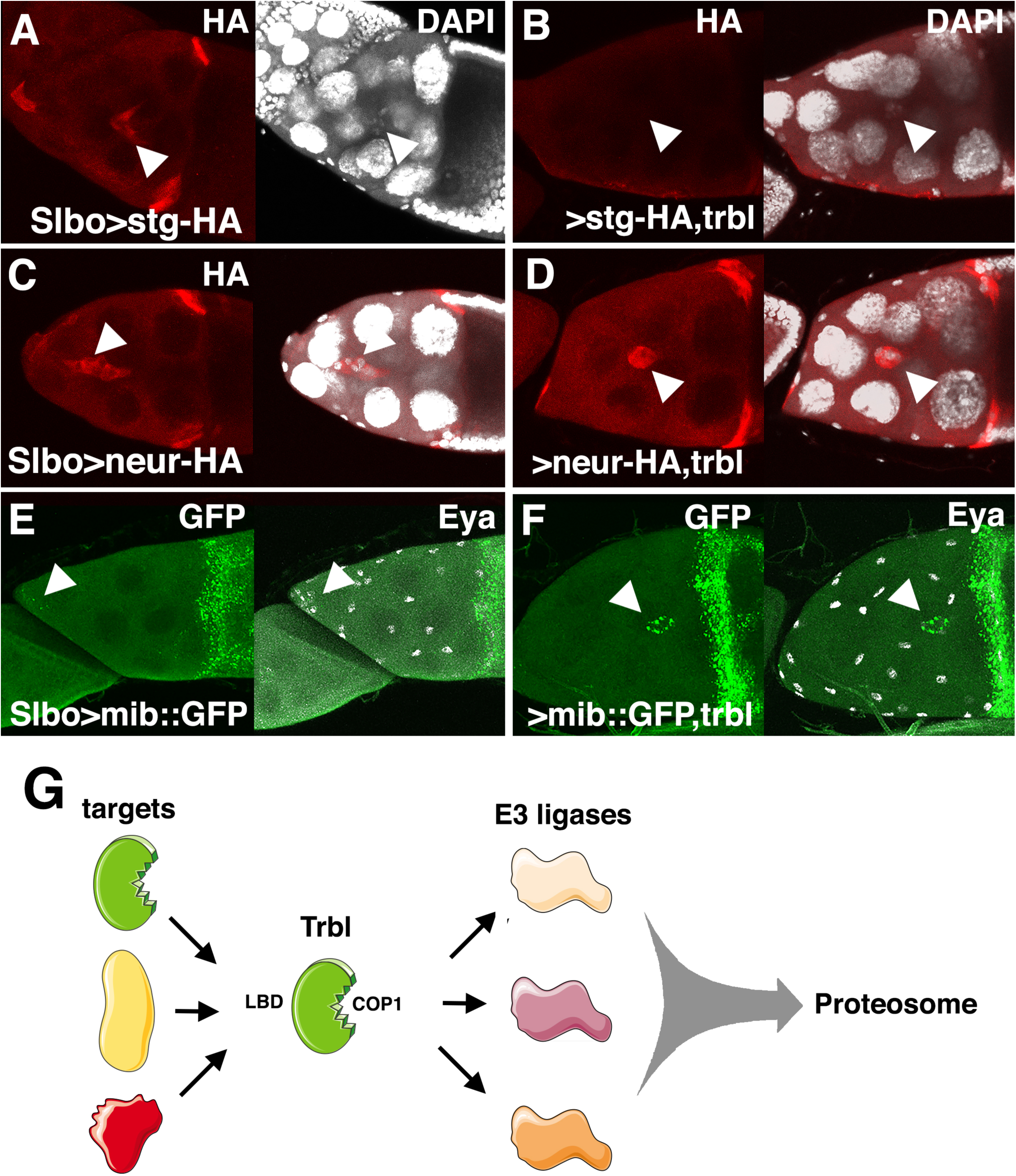
Trbl stabilizes Neuralized in the migrating border cells. A. SlboGAL4 misexpression of UAS-HA-string resulted in intermediate levels of HA staining in the migrating border cells (arrowhead). B. SlboGAL4 misexpression of UAS-HA-string and UAS-trbl resulted in reduced levels of HA staining in the migrating border cells compare to (A), consistent with the ability of Trbl to direct String turnover (arrowhead). C. SlboGAL4 misexpression of UAS-HA-neur resulted in intermediate levels of HA staining in the migrating border cells (arrowhead). D. SlboGAL4 misexpression of UAS-HA-neur and UAS-trbl resulted in increased levels of HA staining in the migrating border cells compare to (C), consistent with the ability of Trbl to stabilize Neur (arrowhead). E. SlboGAL4 misexpression of UAS-Mib1::YFP resulted in intermediate levels of punctate GFP in the migrating border cells (arrowhead). F. SlboGAL4 misexpression of UAS-Mib1::YFP and UAS-trbl resulted in increased GFP levels compared to UAS-Mib1::YFP alone (E), suggesting that Trbl stabilizes Mib1. G. Model for Trbl’s role as an adaptor protein mediating target/E3 ligase interactions via its ligand binding domain (LBD) and COP1 binding site, leading to proteosomal degradation. See text for details.

While on its own SlboGAL4 misexpression of UAS-neur had no effect on border cell migration, Trbl misexpression using the SlboGAL4 driver blocks border cell migration (Rorth et al., 2000; Masoner et al., 2013) so that 54% of egg chambers examined had a complete block to border cell migration and 46% had a partial block (Supplemental Fig. 4E and Supplemental Table 4). Co-misexpression of Trbl and Neur reduced these values to 0% complete block and 40% partial block. Thus, just as Neur can block the Trbl cell size and bristle phenotypes, Neur effectively blocks the Trbl border cell migration defect.

Recently it was demonstrated that mammalian Trib3 binds and stabilizes the E3 ligase Mindbomb, a Neur homolog (Izrailit et al., 2017), so we examined interaction between Trbl and the Drosophila homolog Mindbomb1 (Mib1), a gene with roles complementary to Neur to modulate Notch signaling during development (Lai et al., 2005; Wang and Struhl, 2005). In a yeast two hybrid assay, Trbl interacts with Mib1 with a strength similar to Neur (Fig. 4J), so we used EnGAL4 to misexpress a UAS-mib1 transgene and observed wing vein defects that were suppressed by co-misexpression of UAS-trbl (Fig. 4E,F). Complementarily, Fijiwings revealed that the UAS-mib1 transgene acted as a weak but significant suppressor of the Trbl large cell phenotype (Fig. 4I).

EnGAL4 misexpression of a UAS-Mib1::YFP fusion (Le Borgne et al., 2005) appeared to be more potent than the UAS-mib1 transgene and led to lethality on its own that was effectively suppressed by Trbl co-expression, resulting in a split wing phenotype (Fig. 4G,H). As shown in Fig. 5C, EnGAL4 misexpression of UAS-Mib1::YFP led to strong reduction of Wg expression at the margin, specifically in the posterior compartment. Co-misexpression of Trbl and Mib1::YFP suppressed this effect, restoring Wg expression at the margin and leading to higher Wg levels in cells adjacent to the margin (arrows, Fig. 5D). This expansion of Notch activity in cells adjacent to the margin (marked by arrows in greater detail in Fig. 5E) is consistent with a disruption in patterning at the margin resulting the split wing phenotype observed in the mature wing (Fig. 4F).

While in the wing disk levels of Mib1::YFP were comparable in the presence of absence of Trbl (cf. Fig. 5C,D), in the migrating border cells misexpression of Mib1::YFP resulted in punctate GFP that increased in levels when Trbl was co-misexpressed (Fig. 7E,F). Similar to Neur, Mib1 misexpression had no strong effect on border cell migration on its own but effectively suppressed the Trbl block in border cell migration when co-misexpressed reducing the 54% complete block and 46% partial block associated with Trbl misexpression to 8% complete/0% partial and 0% complete/0% partial following co-misexpression of two different Mib1 transgenes (Supplemental Fig. 4E and Supplemental Table 2). These data suggest that Trbl binds targets like Neur and Mib1 to stabilize their levels and block their activity in the border cells, exactly as we observed at the wing disk margin and in the sensory organ precursors.

## DISCUSSION

### A screen for Trbl targets identifies E3 ligases as suppressors of Tribbles misexpression phenotypes

To better understand how the Tribbles family proteins influence development and disease, we sought novel targets of fly Trbl using a misexpression screen in the Drosophila wing, which is patterned by several conserved signaling pathways affecting tissue and cell size including BMPs, Wnts, Hedgehog and Notch (Beira and Paro, 2016). Misexpression of Trbl in the wing results in a distinct phenotype marked by smaller wing size and larger cells (Fig. 1) due in part by the ability of Trbl to bind the targets Akt and String/Twine-Cdc25 to reduce cell growth and proliferation, respectively (Mata et al., 2000; Farrell and O’Farrell, 2013; Das et al., 2014; Liang et al., 2016b). Consistent with the opposing effects of Stg and Trbl on G2, misexpressing Trbl in the posterior compartment increased CycB levels compared to the anterior compartment while misexpression of Stg decreased CycB levels, and the co-expression of both resulted in similar levels in both compartments (Fig. 2). The sensitivity of the wing system to interactions between Trbl and bona fide targets like Stg and Slbo led us to test its interactions with a set of strains co-expressing UAS-regulated open reading frames (ORF) encoding growth regulating genes affecting wing patterning, size and growth.

Because misexpression approaches can generate artificial phenotypes resulting from ectopic and abnormally high gene levels, we focused on subtle changes in Trbl wing cell and tissue size phenotypes and precisely measured these with Fijiwings, a computer-based tool and from this approach we identified enhancers and suppressors of Trbl (Fig. 3).

We focused initially on several E3 ubiquitin ligase suppressors because the C-terminal tail of mammalian Trib molecules binds the conserved WD40 β-propeller domain of E3 ligases with high affinity (Murphy et al., 2015; Uljon et al., 2016). Trbl suppressed the wing notch phenotype caused by misexpression of the E3 ligase Neuralized (Neur; Fig. 4), and similarly Trbl blocked the split wing phenotype caused by misexpression of Mib1, another E3 ligase with complementary functions to Neuralized (Le Borgne et al., 2005; Wang and Struhl, 2005). Consistent with the ability to suppress both Neur and Mib1 wing notch phenotypes, Trbl co-misexpression with either gene increased levels of the Notch target gene Wg at the wing disk margin (Fig. 5). Trbl binds Neur and Mib1 with similar affinity in yeast two hybrid analysis and notably, suppression of Neur phenotypes by Trbl was not dependent on the conserved RING domain in Neuralized, consistent with the ability of mammalian Trib1 to bind the E3 ligase COP1 when its RING domain is deleted (Uljon et al., 2016).

In the migrating border cells, misexpression of either Neur or Mib1 antagonized the ability of Trbl to block border cell migration (Fig. 7). The ability of Trbl to bind and block the effect of these E3 ligases in the wing and ovary is consistent with the notion put forward by Uljon and colleagues that Trib1 binding to E3 complexes has the potential to redirect the formation of cullin-based proteosomal complexes to other targets (Uljon et al., 2016). Moreover, the identification from our limited wing screen of the E3 ligases Neuralized, Mindbomb1 and Parkin suggests that expanding this screen has the potential to uncover more E3 ligases that use Trbl as a bridge to reach an extended pool of proteosomal degradation targets (Fig. 7H).

### A conserved role for Trbl in mediating E3 ligase-dependent Notch signaling

In Drosophila, the Notch signaling pathway controls cell fate, differentiation, proliferation, apoptosis and stem cell self-renewal (Siebel and Lendahl, 2017). Sensory organ precursor (SOP) cell fate is determined by asymmetric cell division coordinated with localization of Neuralized protein, which simultaneously blocks Notch activation in pIIb and activates Notch signaling in pIIa cells to pattern the socket and shaft (Le Borgne and Schweisguth, 2003). In screens for molecules with roles in sensory organ development, misexpression of Trbl in SOPs resulted in loss of macrochaetes due to the adoption of sheath cell and neuronal cell fates (Abdelilah-Seyfried et al., 2000; Schweisguth, 2015). Detailed examination showed Trbl misexpression transformed pI to pIIb cell fates, with the corresponding loss of the pIIa cell fate, indicating that Trbl exerts a potent block in both Notch signaling and mitosis during the ordered asymmetric divisions required for bristle formation. Our unpublished work showing a role for Trbl and Neur in the mitotic to endocycle switch in the ovarian follicle cells supports the notion that the Trbl/Neur interaction links the cell cycle with Notch activation in at least two tissues (Nauman and Dobens, unpublished).

We show that Trbl/Neur or Trbl/Mib1 co-misexpression can suppress the loss of macrochaetae associated with misexpression of Trbl, Neur or Mib alone (Fig. 6). The connection of Trbl to Notch signaling is significant in light of the association of Tribs 2 and 3 in mammalian systems with Notch-dependent tumor formation (reviewed in Stein et al., 2015). Trib2 was identified as a downregulated gene upon γ-secretase inhibitor treatment of Notch-dependent mouse T cell acute lymphoblastic leukemias cell lines, and the absence of Trib2 in a murine Trib2-knockout model was found to accelerate Notch1-driven leukemias (Liang et al., 2016a; Stein et al., 2016). A correlation between Trib3 and Notch1 expression in lung adenocarcinoma cell lines led to work showing that Notch1 could be downregulated by the knockdown of Trib3 (Zhou et al., 2013). High-throughput kinase inhibitor screens identified Trib3 as a master regulator of Notch through the MAPK-ERK and TGFβ pathways in breast cancer (Izrailit et al., 2013).

Recently, Trib3 has been shown to promote Notch-induced tumor growth by acting as an adaptor between the deubiquitinase USP9x and the E3 ligase Mindbomb (Mib), to efficiently stabilize Mib levels and promote ligand-dependent Notch activation (Izrailit et al., 2017). In this light, our results showing that Trbl can stabilize levels of Neur and Drosophila Mib1 suggests that its role as a scaffold adaptor protein is conserved with the mammalian Notch pathway. Because the mechanism of Notch receptor activation is simplified in flies, with one Notch receptor and the two ligands Delta and Serrate (Schwanbeck and Just, 2011; Guruharsha et al., 2012), going forward the Drosophila system will offer an excellent place to combine loss-of-function with epistatic tests to better understand how Trbl modulates the activity of E3 ligases to regulate the direction and strength of Notch signaling.

## MATERIALS AND METHODS

### Drosophila strains

Stocks used were: (1) *en2.4-GAL4* (EngrailedGAL4)*/CyO;Trbl/TM3*, (2) UAS-ORF collection listed in the Supplemental table 1 (Bischof et al., 2013), (3) *UAS-HisMyc-mib A1-3A* (on III), a gift from Dr. Eric Lai Memorial Sloan Kettering Cancer Center (Lai et al., 2005), (4) *UAS-Mib1::YFP 3-1* (gift from Dr. Francois Schweisguth, Institut Pasteur), (5) *UAS-slbo* (gift from Dr. Pernille Rorth), (6) *w P{w*(+mW.hs)*=GawB}neur*(GAL4-A101) *Kg*(V)*/TM3, Sb*(1) (NeurGAL4) (Rulifson et al., 2002), (7) *M{UAS-neur.ORF.3xHA}ZH-86Fb* (UAS-neur 3xHA), (8) *P{UAS-FLAG-trbl.WT}* (Masoner et al., 2013), (9) *y*(1) *w*(67c23); *P{w*(+mC)*=UAS-neur.* Δ*RING::EGFP}8A* (NeurΔRING), (10) *y*(1) *v*(1); *P{y*(+t7.7) *v*(+t1.8)*=TRiP.JF02048}attP2* (Neur RNAi).

### Wing photomicrography and Fijiwings analysis

Wings were dissected and prepared from female flies reared at 30OC (for increased GAL4 activity) as described in (Dobens and Dobens, 2013). Wing compartment size was measured in kilopixels and trichome density was measured in units per kilo pixel using Fijiwings software (https://sourceforge.net/projects/fijiwings/). Fijiwings was also used to generate heat map images in Fig. 1.

In Fig. 3, the trichome density ratio was calculated by dividing the trichome density measurement of the first anterior intervein region by the trichome density measurement of the third posterior intervein region. Each point on this plot is the average of 3-4 trichome density ratios. The area ratio was calculated by dividing the area measurement in pixels of the first anterior intervein region by the area measurement of the third posterior intervein region. In Supplemental Fig. 2, the average trichome density ratios and average area ratios were plotted against each other to examine how misexpression of Trbl and the UAS-ORFs influenced these factors in a combined manner. Log scales were used on the X and Y axes to better spread data points. In Fig. 4, the trichome density measurement of the third posterior intervein region was divided by the trichome density measurement of the first anterior intervein region. The ratio calculated for Fig. 4 is reversed compared to the ratios calculated for Fig. 3 so a higher ratio will correlate with a greater number of trichomes present in the wing; the error bars represent one standard deviation. Statistical analysis was performed with Graph Pad Prism (v 6.0) and all data are presented as mean ± SD and analyzed using unpaired, 2-tailed Student’s t test or ANOVA followed by post hoc analysis of significance (Tukey’s test). A P-value of less than 0.05 is considered significant. GraphPad Prism (v 6.0) was used to generate the graphs and perform statistical tests.

### Yeast two-hybrid interaction screen

The ProQuest Two-Hybrid System (Invitrogen) was used to perform yeast two-hybrid interactions between Tribbles and the candidate genes. *Saccharomyces cerevisiae* MaV203 cells were transformed according to the kit manual and grown on SC media lacking tryptophan and leucine to ensure presence of bait and prey plasmids. To test for interaction, cells were spotted onto SC-Leu-Trp-His+3-Amino-1,2,4-triazole (3-AT) (at 0mM, 10mM, 25mM, and 100mM concentrations) plates, and grown at 30C for 5 days. Construction of FLAGTrbl bait and prey plasmids has been previously described (Masoner et al., 2013). Bait and prey plasmids for Neur and Mib1 were constructed by PCR amplifying genes from the cDNAs LD45505 and SD05267 (from the Drosophila Genomics Resource Center), using primers incorporating flanking *attB1* and *attB2* sites. The forward and reverse primers for these genes are as follows:

For Neur 5’- **GGGGACAAGTTTGTACAAAAAAGCAGGCT**CGATGGGTCTATCGGATATAC CAGCC -3’, 5’- **GGGGACCACTTTGTACAAGAAAGCTGGGT**GTCACTACGTGGTGTAGGTGC GGATGACG -3’; for Mib1 5’- **GGGGACAAGTTTGTACAAAAAAGCAGGCTCG**ATGTCTTGTGCGGCCACCC TCTCG

-3’, 5’- **GGGGACCACTTTGTACAAGAAAGCTGGGTCCTA**TCAGAAGAGCAGGATGC GTTTTTC -3’ (the *attB1* and *attB2* sequences in bold). The PCR products were cloned into the donor vector pDONR-21, and then into the bait vector, pDEST32 and prey vector, pDEST22. All constructs were clones using the Gateway Cloning System (Invitrogen) and clones were confirmed by sequencing. The positive control included in the kit was generated by transforming yeast with pEXP32/Krev1 and pEXP22/RalGDS-wt plasmids, and the negative control consisted of pEXP32/Krev1 and pEXP22/RalGSD-m2 plasmids.

### Immunostaining

Tissue was stained with the following primary antibodies: 1:1000 mouse or rabbit anti HA (Sigma and Cell Signaling Technologies, respectively); 1:200 mouse anti-Eya (Developmental Studies Hybridoma Bank, DHSB), 1:200 anti-Wg (DHSB); 1:200 anti-Dlg (DHSB); 1:1000 rabbit anti-GFP (Sigma), 1:1000 rabbit anti-Bgal (DSHB), 1:200 mouse anti-Wingless (DSHB). Secondary antibodies used were: AlexaFluor 1:200 goat anti-rabbit 594, 1:200 goat anti-mouse 488, 1:200 goat anti-mouse 594, 1:200 goat anti-chicken 594 (Invitrogen). Samples were also stained with DAPI (Roche). Images were collected on an Olympus Fluoview 300 Confocal Microscope or Nikon 90i microscope with Optigrid paddle and Metamorph Image acquisition software. Images were processed using ImageJ and Photoshop Elements 2018.

### Notum images

Flies were sorted and frozen at −20C for 5 minutes and bristles were photographed under diffuse light using a Canon EOS Rebel T5 attached to an Olympus SZ-12 dissecting microscope, and Canon EOS Utility software (version 2.14.31). To create high-quality images, several images were taken at various focal points and then combined using Helicon Focus software (Helicon Soft).

### Statement on reagent and data availability

The authors confirm that the data supporting the findings of this study are available within the article and its supplementary materials.

## ACKNOWLEDGEMENTS

We would like to thank Dr. Eric Lai and Dr. Francois Schweisguth who generously sent Mib1 flies. We thank Dr. Erika Geisbrecht and members of the lab for helpful discussions, especially Jorgues Martinez. We thank Samantha Schwartz and Samuel Gutierrez for assistance in producing Supplemental Fig. 3 and Laura Dobens for assistance in the production of Supplemental Fig. 4. Funding support came from the NSF (IOS-1456023), NIH (NIH R21 CA197317) to LLD; UMKC School of Graduate Studies (Graduate Student Research Award Program 2014-2017) and the UMKC Women’s Council (Graduate Assistance Fund, 2014-2016) to AS and CN; the UMKC ADVANCER (Academic Development Via Applied and Cutting-Edge Research) award to BH.

## SUPPLEMENTAL FIGURES

**Supplemental Figure 1. Opposing effects of Trbl and Stg on Cyclin B stability**

A. EnGAL4 misexpressing UAS-Trbl in posterior wing compartment led to subtly increased levels of RFP-CycB1–266, the fusion of the CycB degron to RFP, and decreased levels of the GFP-E2F11–230 fusion protein fusing the E2F degron to GFP, consistent with posterior cells blocked in G2.

B. EnGAL4 misexpressing UAS-Stg in posterior wing compartment led to decreased levels of RFP-CycB1–266 with no strong effect on GFP-E2F11–230 fusion proteins, consistent with its ability to promote the G2/M transition to G1.

**Supplemental Figure 2. Comparison of trichome density ratio to area ratio** Comparison of the trichome density ratio to the area ratios of all UAS-ORFs co-misexpressed with Trbl by the EnGAL4 driver. The continuum from WT (purple) to Trbl (red) is shown highlighted as a red line.

**Supplemental Figure 3. Trbl misexpression and and Neur knockdown affect bristle patterning.**

A. WT notal bristle pattern.

B. NeurGAL4 driving UAS-GFP expression in mature notum.

C. NeurGAL4 driving misexpression of an RNAi knockdown construct for neuralized results in loss of macrochaetae and disorganization of microchaetae.

D. NeurGAL4 driving misexpression of UAS-Trbl results in loss of macrochaetae bristles and disorganization and loss of microchaetae.

**Supplemental Figure 4. Trbl stabilizes Neuralized in the migrating border cells under the influence of the enhancer trap driver GAL4-Hsp70.PB**

A. GAL4-Hsp70.PB misexpression of UAS-HA-string resulted in intermediate levels of HA staining in the migrating border cells. Note that for A-D, flies were reared at 25oC and not subjected to heat shock; under these conditions, border cell expression represents enhancer trap activity of this driver.

B. GAL4-Hsp70.PB misexpression of UAS-HA-string and UAS-trbl resulted in reduced levels of HA staining in the migrating border cells compare to (A), consistent with the ability of Trbl to direct String turnover.

C. GAL4-Hsp70.PB misexpression of UAS-HA-neur resulted in intermediate levels of HA staining in the migrating border cells.

D. GAL4-Hsp70.PB misexpression of UAS-HA-neur and UAS-trbl resulted in increased levels of HA staining in the migrating border cells compare to (C), consistent with the ability of Trbl to stabilize Neur.

E. Summary of border cell migration defects from transgene misexpression by SlboGAL4 driver.

**Supplemental Table 1. Fijiwings data**

SupplementalTable1.xlsx

**Supplemental Table 2. Area interactors**

SupplementalTable2.xlsx

**Supplemental Table 3. Trichome density interactors**

SupplementalTable3.xlsx

**Supplemental Table 4. Trichome density interactors**

SupplementalTable4.docx

